# Shallow Recurrent Decoders for Neural and Behavioral Dynamics

**DOI:** 10.1101/2025.06.20.660769

**Authors:** Amy S. Rude, J. Nathan Kutz

## Abstract

Machine learning algorithms are affording new opportunities for building bio-inspired and data-driven models characterizing neural activity. Critical to understanding decision making and behavior is quantifying the relationship between the activity of neuronal population codes and individual neurons. We leverage a SHallow REcurrent Decoding (SHRED) architecture for mapping the dynamics of population codes to individual neurons and other proxy measures of neural activity and behavior. SHRED is a robust and flexible sensing strategy which allows for decoding the diversity of neural measurements with only a few sensor measurements. Thus estimates of whole brain activity, behavior and individual neurons can be constructed with only a few neural time-series recordings. This opens up the potential for using non-invasive, or minimally invasive, measurements for estimating difficult to achieve, or invasive, large scale brain and neural recordings. SHRED is constructed from a temporal sequence model, which encodes the temporal dynamics of limited sensor data in multiple scenarios, and a shallow decoder, which reconstructs the corresponding high-dimensional neuronal and/or behavioral states. We demonstrate the capabilities of the method on a number of model organisms including *C. elegans*, mouse, zebrafish, and human biolocomotion.

## 1 Introduction

Advances in neuroscience over the past decades have revealed that the nervous systems leverages dimensionality reduction when it encodes behavior and decision making, collapsing the highdimensional representation of the stimulus environment into a much lower representation of motor command or output behaviors [1–4]. Thus instead of relying on individual neurons, one can consider the dynamic encoding space, or low-dimensional manifold [5], of the brain for decision making and cognitive capabilities. This dimensionality reduction reflects the brain’s ability to efficiently organize complex computations, enabling insights into how neural circuits generate behavior and cognition [6, 7]. A second critical observation is that sparsity plays a key role in encoding and decoding information in the network [8–10]. These two observations are the basis of our proposed *SHallow REcurrent Decoder* (SHRED); a deep learning architecture that allows for an efficient and robust mathematical framework connecting the activity of individual (sparse) neurons to population codes and behavioral measurements. Thus the monitoring of a small number of neurons or their proxies is capable of reconstructing brain-wide activity or proxy behavioral outcomes, as we demonstrate in the nematode Caenorhabditis elegans (*C. elegans*), the mouse, zebrafish and in human biolocomotion.

High-dimensional networked biological systems are ubiquitous and characterized by a large connectivity graph whose structure determines how the system operates as a whole [11, 12]. Typically the connectivity is so complex that the functionality, control and robustness of the network of interest is impossible to characterize using currently available methods. Moreover, with few exceptions, underlying nonlinearities impair our ability to construct analytically tractable solutions, forcing one to rely on experiments and/or modern high-performance computing to study a given system. But as noted, advances in experimental and computational neuroscience over the past decades have revealed two critical, and seemingly ubiquitous, phenomena: (i) that meaningful input/output of signals in high-dimensional networks are encoded in low-dimensional patterns of dynamic activity called population codes [1–4], and (ii) that sparsity is fundamental in encoding and decoding information in the network [8–10]. A full understanding of the computational process encoded throughout a nervous system must reconcile these two key observations that transforms sensory input into decision making and motor representations.

Originally developed as a sensing strategy, SHRED [13] is a promising deep learning architecture with impressive reconstruction and forecasting results in the low-data limit [14–18]. Importantly, SHRED’s basic architecture connects sparse sampling of signals to population codes for applications in neuroscience. SHRED leverages the mathematical theory of Takens’ embedding theorem [19] to trade population code information for temporal trajectory information of a single neuron across time. Previous work has shown SHRED can achieve excellent performance in sensing and reduced order modeling examples ranging from weather and atmospheric forecasting [13], plasma physics [14], nuclear reactors [16], and turbulent flow reconstructions [17]. Theoretically rooted in the classic PDE method of separation of variables [13, 17], the decoding-only strategy of SHRED circumvents the computation of inverse pairs, i.e. an encoder and the corresponding decoder. It has been well-known for decades that the computation of the inverse of a matrix is highly unstable and ill-posed [20, 21]. By decoding only, SHRED avoids this problem and learns a single embedding without the corresponding inversion.

SHRED is leveraged on a number of toy models in neuroscience, including the nematode Caenorhabditis elegans (*C. elegans*), the mouse, zebrafish, and human biolocomotion. In each case, SHRED is shown to be robust and accurate in approximating population codes using only a limited number of neurons. Figure 1 details the basic architecture of the neural network, where the latent space of a temporal encoder is used to reconstruct full activity across neurons in the

**Figure 1.**
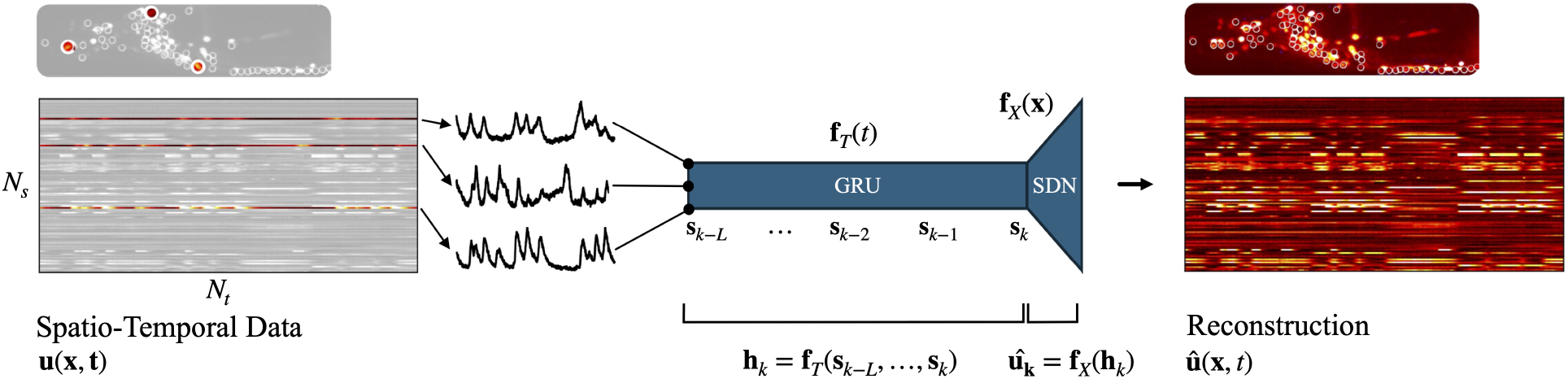
Diagram of *SHallow REcurrent Decoder* (SHRED) architecture. Time-series data from three neurons 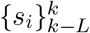 is fed into a *Gated Recurrent Unit* (GRU). The output from GRU **h**_**k**_ is the input for the *Shallow Decoder Network* (SDN), mapping back to the full state space. *C. elegan* neuron image (top) is adapted from [27].

*C. elegans*. An extensive study is made for each model system showing that the architecture can be used to critically reconstruct population codes from only a highly limited number of individual neuron recordings. In the case of human biolocomotion, it is used instead to estimate gait motion using only a single sensor (smart watch IMU). The architecture is thus promising for extracting critical information on decision making and motor commands with only limited, and potentially non-invasive, measurements. This provides a valuable potential reduction to practice for digital healthcare for the management of functional neurological disorders by encoding critical metrics of cognition through proxy measurements encoded through SHRED.

## 2 Mathematical Background for SHRED

SHRED consists of a combination of recurrence and decoding, with the former operation capturing the temporal behavior of limited sensor data and the latter operation performing a spatial upscaling to recover the corresponding high-dimensional state. A similar split in spatial and temporal components is taken into account by the well-known method of separation of variables for solving linear PDEs [22], which assumes that the solution has the form *u*(**x**, *t*) = *T* (*t*)*X*(**x**). Indeed, it has been shown that in the limit of linear dynamics with constant coefficients, the SHRED architecture has a closed form analytic solution [13, 17]. Indeed, in the linear limit the analytic solution is given by the dynamic mode decomposition (DMD) [23, 24]. Of note is that the architecture can be used to produce estimates for uncertainty quantification [25] along with multiscale features [26].

Thus SHRED is a generalization of separation of variables with nonlinear (neural network) approximations. A compact representation of the SHRED algorithm, as shown in Fig. 1 is as two neural networks jointly trained to the form

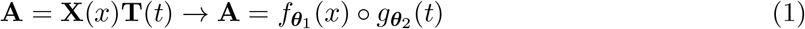

where **A** is a data matrix whose columns are time samples and whose rows are the full state space, *g****θ*** _2_ (*t*) is trained largely to encode sequential (time) data and 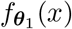 is largely responsible for mapping from the latent space of the sequential encoding to the full state-space. Of course, the time and space encoded are not independent as they are jointly trained and the decoder is dependent on the time history of the sequence model.

To be more precise, as a deep learning architecture, SHRED aims to reconstruct high-dimensional neuronal data from *N*_*s*_ individual neural recordings. Instead of learning the *one-shot* reconstruction map 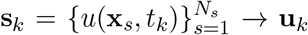 for *k* = 1, …, *N*_*t*_ – as considered by, e.g., [28–31] – we take advantage of the temporal history of the individual neuron values. Specifically, a recurrent neural network **f**_*T*_ encodes the neuron measurements over a time window of length *L ≤ N*_*t*_ into a latent representation of dimension *N*_*l*_, that is

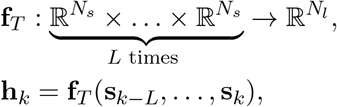

where the hyperparameter *L* identifies the number of lags considered by SHRED-ROM, and may be selected according to the problem-specific evolution rate or periodicity. The high-dimensional state is then approximated through a decoder **f**_*X*_, which performs a nonlinear projection of the latent representation onto the state space

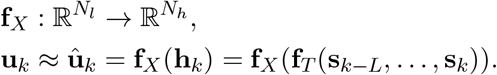

In this work, we consider a *gated recurrent unit* (GRU) network [32] to model the temporal dependency of sensor data, and a *shallow decoder network* (SDN) as latent-to-state map. SHRED is a modular framework, allowing for various forms of temporal encoding and spatial decoding, including convolutional neural networks [33], LSTM [34], echo state networks [35], transformers [36] and variational autoencoders [37]. Importantly, the decoding only strategy circumvents learning an encoder-decoder pair [38] which leads to numerical stiffness [20, 21].

Unless noted otherwise, the data used for training SHRED is accomplished with an 80:20 train/test split. Each data set is unique, and amount of data in the 80:20 split can differ significantly between the *C. elegans*, mouse, and zebrafish. If a different train/test split is used, it is explicitly outlined below. For details of training and access to the code used to produce the results, see Sec. 5. Table 1 gives a comprehensive summary of all mean-squared error results for Sections 3 A, B and C.

**Table 1:**
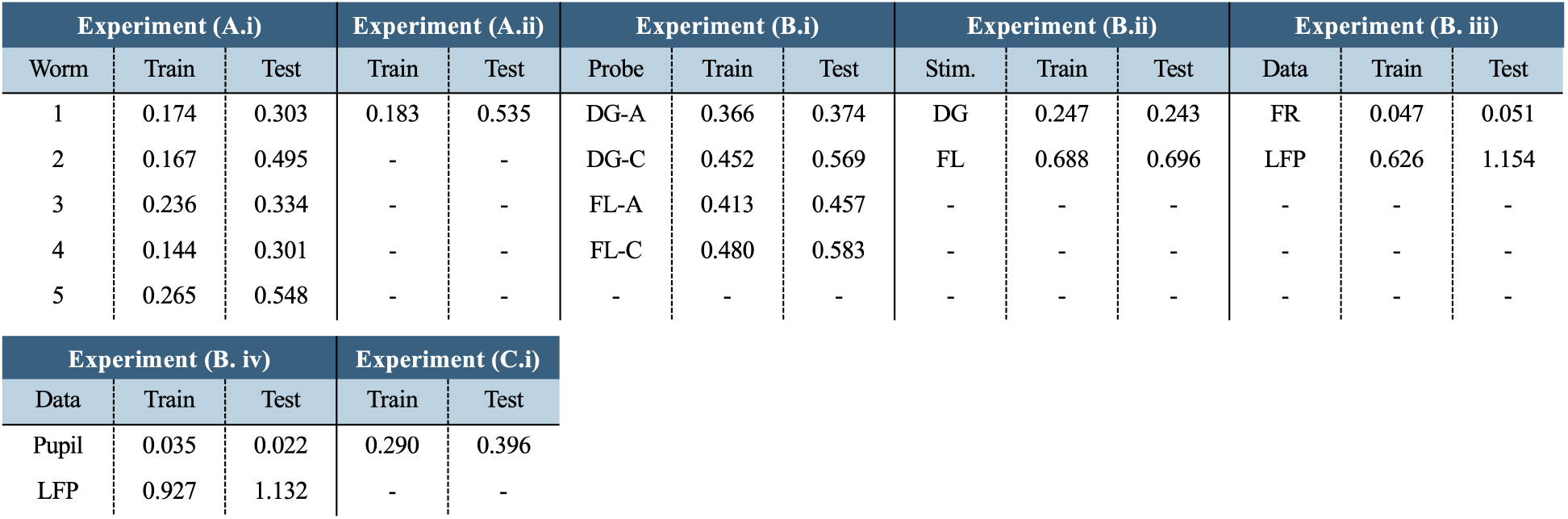
Mean Square Error (MSE) for all experiments presented in Sections 3.1, 3.2, and 3.3. DG - drifting grating stimulus. FL - flash stimulus. FR - firing rate. LFP - local field potential.

| Experiment (A.i) |  |  | Experiment (A.ii) |  | Experiment (B.i) |  |  | Experiment (B.ii) |  |  | Experiment (B. iii) |  |  |
| --- | --- | --- | --- | --- | --- | --- | --- | --- | --- | --- | --- | --- | --- |
| Worm | Train | Test | Train | Test | Probe | Train | Test | Stim. | Train | Test | Data | Train | Test |
| 1 | 0.174 | 0.303 | 0.183 | 0.535 | DG-A | 0.366 | 0.374 | DG | 0.247 | 0.243 | FR | 0.047 | 0.051 |
| 2 | 0.167 | 0.495 | - | - | DG-C | 0.452 | 0.569 | FL | 0.688 | 0.696 | LFP | 0.626 | 1.154 |
| 3 | 0.236 | 0.334 | - | - | FL-A | 0.413 | 0.457 | - | - | - | - | - | - |
| 4 | 0.144 | 0.301 | - | - | FL-C | 0.480 | 0.583 | - | - | - | - | - | - |
| 5 | 0.265 | 0.548 | - | - | - | - | - | - | - | - | - | - | - |

| Experiment (B. iv) |  |  | Experiment (C.i) |  |
| --- | --- | --- | --- | --- |
| Data | Train | Test | Train | Test |
| Pupil | 0.035 | 0.022 | 0.290 | 0.396 |
| LFP | 0.927 | 1.132 | - | - |

## 3 Model Systems

The SHRED architecture maps limited, time-series sensor measurements to the full, high-dimensional neural or behavior data. In what follows, we show how measuring only a limited number of neurons or sensors can estimate the full state space in four different model systems. This demonstrates the capability of the algorithm for handling challenging data.

### 3.1 *C. elegans* nematode

The nematode (*C. elegans*) is a perfect model organism to consider in the context of neuro-sensory integration as it is comprised of only 302 sensory, motor and inter-neurons whose electro-physical connections (i.e. its connectome) are known and stereotyped [39,40]. In addition, it possesses only a small number of sensory neurons, often linked to specific stimuli [41]; neural activity can be imaged in real time; and its range of behavioral responses, during chemotaxis for instance, are varied yet limited, approximately confined to five observable motor states that are related to forward and backward crawling, omega turns, head sweeps and brief pause states. Thus it is reasonable to posit a complete model of its neuro-sensory integration capabilities. Aiding in this effort is the near-complete connectivity data for the gap junctions and chemical synapses connecting the sensory neurons to the inter- and motor-neurons [42]. Importantly, modern tracking microscopes provide rich data sets that enable one to not only record brain-wide activity, but also track the movements and postures of a worm as they explore their environments. With access to a complete record of sensory neuron activity across the sensory periphery as well as a complete record of command motor neuron activity and locomotor behavior, we are well poised to understand how decision-making circuits collapse sensory representations into the most basic commands that shape behavior by regulating muscle activity. Recent experiments in behaving animals retain proprioceptive feedback features which strongly shape computational pathways in sensorimotor circuits [43, 44].

We obtained whole-brain calcium imaging recordings of immobilized nematodes (*C. elegans*) from a publicly available dataset provided in [27]. This dataset contains data for five worms with recordings from 107 - 131 identified neurons in an environmentally controlled condition. Recordings were made for a duration of roughly 18 minutes at an average rate of 2.85 Hz. Since bleaching causes damping of *Ca*^2+^ imaging signals over time, we truncated the data to only include the first half of each session [27]. In the following, we apply SHRED to this dataset. We reconstructed neural activity for each individual worm in Section 3.1.1. SHRED was also applied across worms to predict neural activity in a worm that was hidden during training in Section 3.1.2. These results demonstrate the capabilities of SHRED to decode population-wide dynamics from sparse neural measurements.

#### 3.1.1 SHRED reconstruction of *C. elegan* Neural Activity

We reconstructed whole population activity for all five worms using SHRED. As shown in the carpet plots in Figure 2, some neurons remained inactive for the entire duration of the recording. To ensure that our model performance was not negatively impacted by the selection of these inactive neurons, we created subgroups of *High Variance* (HV) and *Low Variance* (LV) neurons. Details of this split can be found in Section 3.1.3. To perform the reconstruction, we randomly selected three neurons from the HV group and the first 1500 timepoints (approximately 8.5 minutes) from each worm was split 80-10-10 (train-test-validation). With a lag *L* = 100 the model was trained for 200 epochs. The corresponding Mean Square Error (MSE) was calculated for both the training and testing datasets.

**Figure 2.**
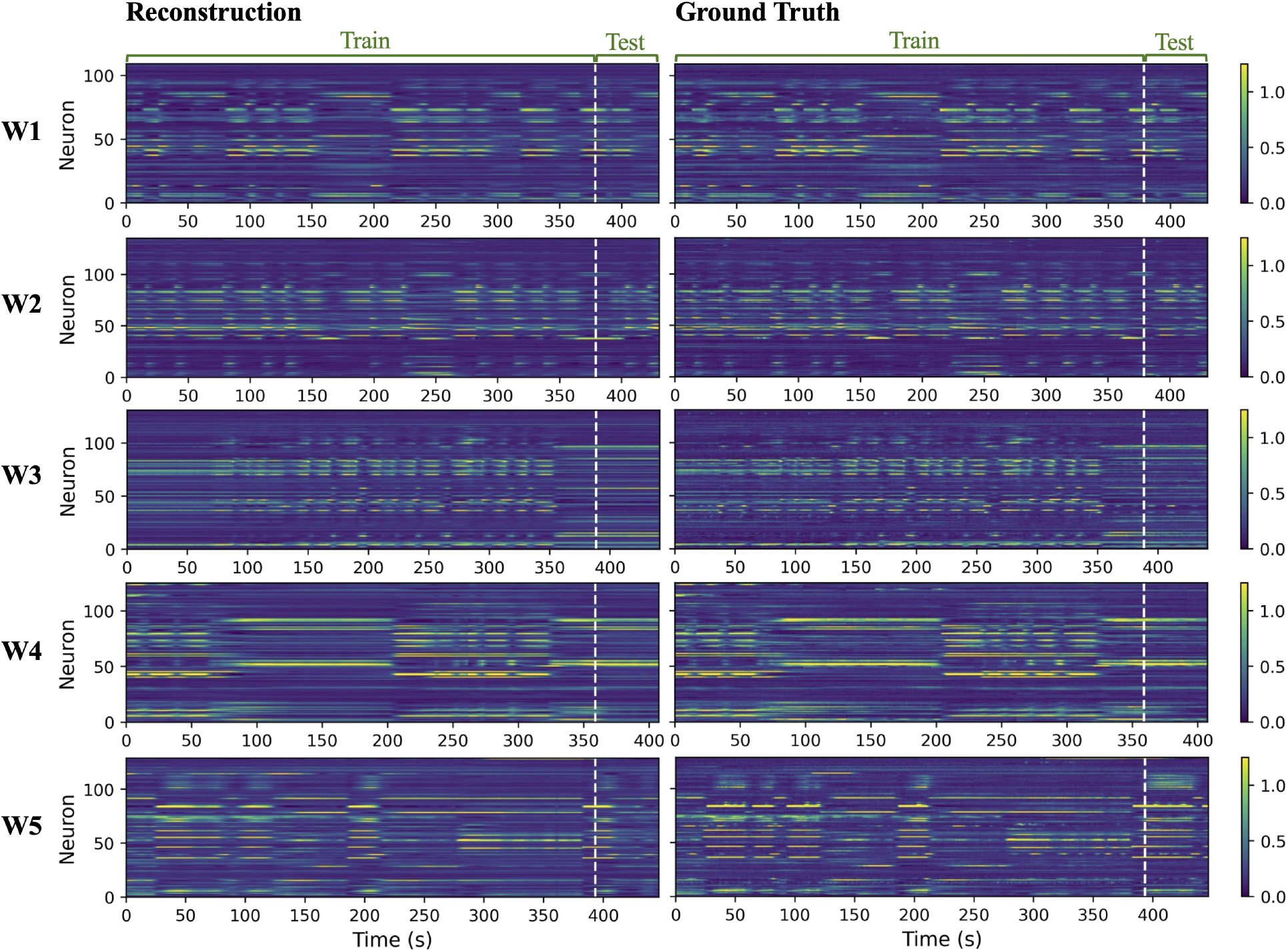
SHRED can accurately reconstruct population-wide activity from only three neurons. SHRED reconstruction (left) of neural activity (right) for all five worms. General activity patterns are preserved in the reconstruction of both training and testing datasets.

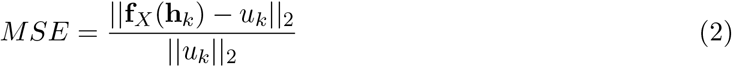

SHRED does a remarkable job learning spatial and temporal patterns in data. To the naked eye, the reconstructions presented in Figure 2 are near identical to the original data. Activity traces are reconstructed in a temporally precise fashion —the peaks and troughs in the activity of each neuron are well-preserved. Even for the withheld test dataset, the model performs remarkably well with the exception of Worm 5 (W5). Several neurons exhibit heightened activity throughout the duration of the test data. This pattern in activity is sustained for a shorter duration in the reconstruction, resulting in a higher error.

The train and test MSE are presented in Table 1(A.i). The errors are slightly higher than what we would have expected from the carpet plots presented in Figure 2. This discrepancy is best explained when looking at individual activity traces. Figure 3 displays in grey, the activity of neuron SMDVR, a cholinergic motor neuron, from W5. This trace is compared to the SHRED reconstruction shown in red and a filtered trace obtained from a low-pass Butterworth filter with a highcut frequency of 0.01 Hz in purple. As shown in this figure, general spikes in activity of SMDVR are maintained in both the SHRED reconstruction and the filtered data. However the magnitude of the fluctuations differ for both the reconstructed and filtered trace, resulting in a higher overall error. Similar to the MSE for test dataset of W5, the filtered test dataset has a MSE of 0.392.

**Figure 3.**
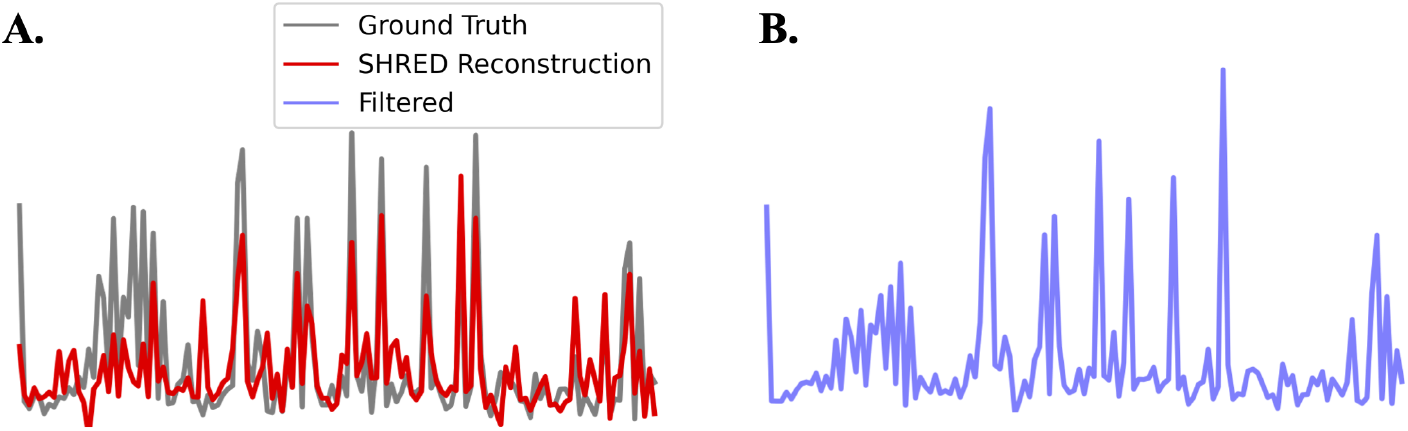
Activity trace for SMDVR neuron. **(A)** - SHRED reconstruction of SMDVR activity (red) overlayed on ground truth (grey). **(B)** - Filtered activity trace. Axis scaling is consistent between the three traces.

#### 3.1.2 Cross-Worm Predictions using SHRED

So far, SHRED has shown remarkable ability for reconstructing neural activity from a select number of neurons for a single individual. In this task, we leveraged the conserved patterns in activity of *C. elegans* to reconstruct neural activity across all five worms. For each worm, we selected the first 1000 time points. Only 45 neurons were recorded from across all 5 worms. Therefore, only these neurons were selected to be included in this trial . The training data was defined to be the first 3000 time points which corresponds to the neural activity of the first three worms. The validation data was defined to be the next 1000 time points corresponding to the neural data of worm four. The test data was defined to be the last 1000 time points which corresponds to the neural data of worm five. Reconstruction of the test dataset only includes the first 800 time points to account for the lag (L) of 200. We randomly selected three neurons from the *High Variance* group for the reconstruction. Results are presented in Figure 4.

**Figure 4.**
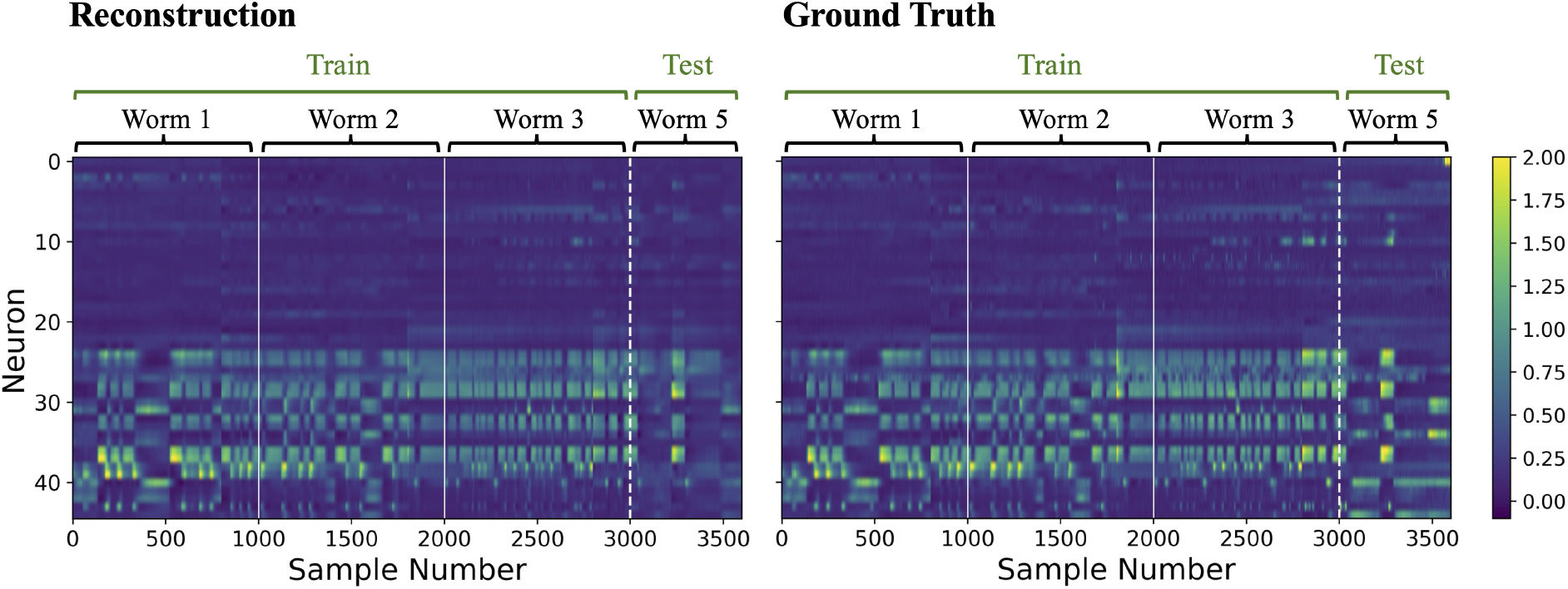
SHRED can generalize across worms. A SHRED model trained to reconstruct neural activity of Worms 1-3 from three *High Variance* neurons can extrapolate to reconstruct activity for Worm 5.

The training error and test error are in Table 1(A.ii). As expected, since the test set is extrapolating to a new worm not included in the training, the test error is significantly higher. However, visually, the general trends in activity are still maintained for the test dataset (Figure 4). With the ability to extrapolate beyond individuals used for training, SHRED can clearly learn distinct spatio-temporal patterns without overfitting to the training data.

#### 3.1.3 Sensitivity Analysis of SHRED Performance

We also conducted some preliminary tests to study the sensitivity of the SHRED model to 1) the activity of the neuron selected, 2) the number of neurons selected and 3) the data split. For these series of experiments, a limited number of neurons were selected and SHRED was trained to predict population activity. Only the first half of the time-series data for the first *C. elegan* (W1) was used to simplify analysis.

Out of the 109 neurons recorded from W1, approximately half exhibited minimal fluctuations in activity over the entire recording period (Figure 5 A-B). This suggests that a substantial proportion of neurons remained inactive throughout the trial thus the selection of these neurons for population-wide predictions with SHRED may result in sub-optimal results. Standard deviations across time for each of the neurons were calculated and neurons with activity levels above the mean were classified as the *High Variance* (HV) group and those below were classified as the *Low Variance* (LV) group. To test the effect of neuronal variability on SHRED performance, three neurons were randomly selected from each of the following groups: HV, LV, and the full population (FP). SHRED was trained to reconstruct the entire population activity. The corresponding train and test Mean Squared Error (MSE) were calculated. We repeated these trials for 25 independent runs.

**Figure 5.**
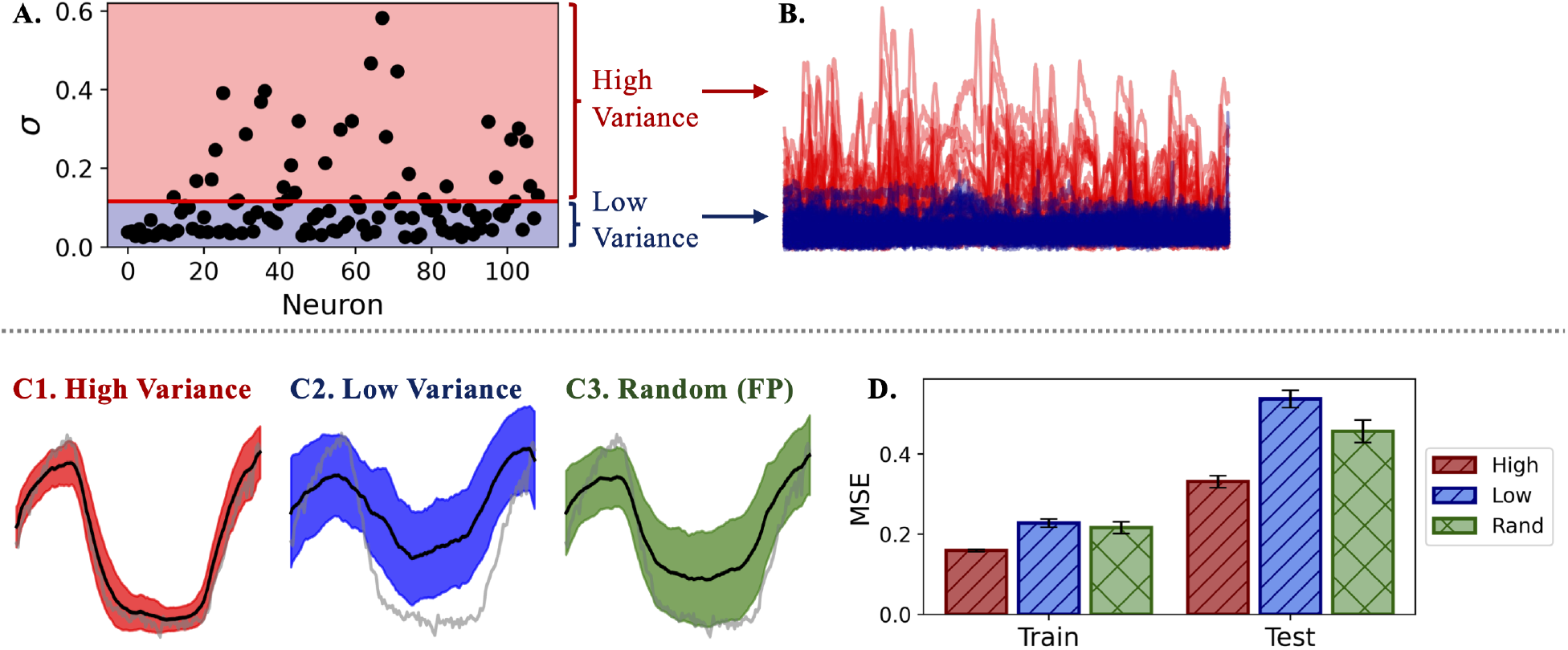
Selecting neurons with highly variable activity improves SHRED performance. **(A)** Scatterplot of 109 neurons (black dots) showing variability in activity, measured as the standard deviation across 1,568 time points. The red line indicates the mean standard deviation. Neurons with values above the average were classified as the *High Variance* (HV) group and those below were classified as the *Low Variance* (LV) group. **(B)** Activity traces from HV neurons (red) and LV neurons (blue) illustrating differences in signal variability. **(C)** A randomly selected neuron trace (grey) and average SHRED reconstruction (black) from 25 trials. Shaded region indicates *±*1 SD. **(D)** Average train and test Mean Squared Error (MSE) for 25 trials using 3 neurons sampled from the HV, LV, or full population (rand).

Randomly selecting from the HV group yielded the best overall performance. Across 25 independent runs, the averaged prediction for the HV condition (Train: 0.159*±*0.003 Test: 0.331*±*0.015) most closely matched the ground truth and exhibited lower variability in performance compared to the LV (Train: 0.228 *±* 0.010 Test: 0.538 *±* 0.022) and FP conditions (Train: 0.217 *±* 0.015 Test: 0.457 *±* 0.028) as shown in Figure 5 C - D. While randomly selecting from the entire population resulted in lower overall MSE compared to the LV condition, there was greater variability in accuracy across individual runs.

Previous applications of SHRED —such as in sea surface temperature, turbulent fluid flow, and magnetoencephalography (MEG) recordings— have demonstrated that as few as three sensors are sufficient to reconstruct whole-system dynamics [45]. To support these results, we then tested the effect of the number of neurons selected on the reconstruction of this immobilized *C. elegan* neural dataset. A total of 15 independent runs were conducted for trials selecting from one, up to five neurons. Train and test MSE were calculated and averaged across trials.

As expected, three neurons are sufficient for accurate reconstruction of the entire population dynamics using SHRED. As shown in Figure 6, the prediction error plateaus after around 3 neurons. There is no substantial additional benefit when including more than three neurons. Analogous to how three sensors are sufficient to localize objects in two dimensions, reconstruction of spatiotemporal data also seems to improve with the inclusion of three sensors.

**Figure 6.**
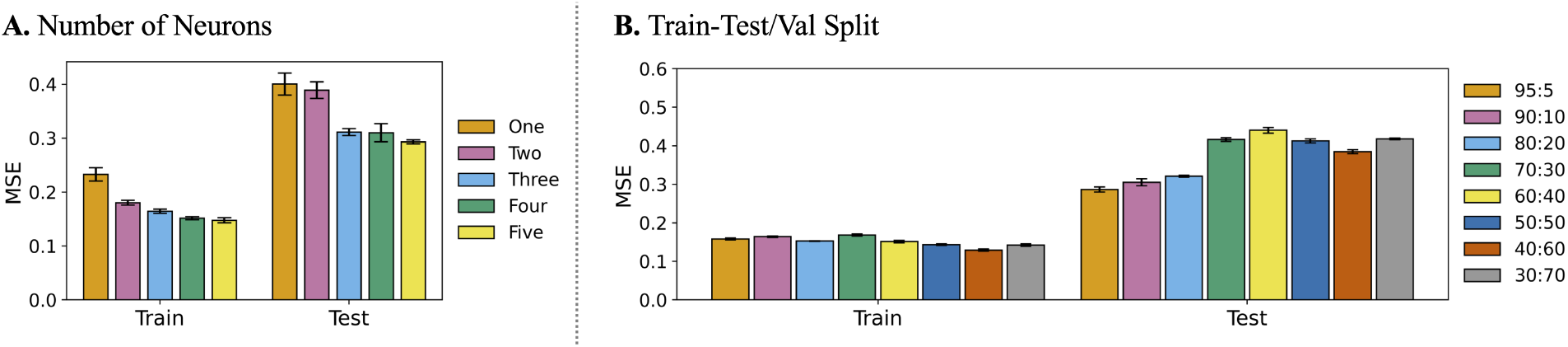
Number of neurons and data split influence SHRED performance. **(A)** Train and test MSE decrease with neuron count but plateaus after three neurons. **(B)** The data split primarily affects reconstruction accuracy on the test dataset, with minimal impact on the training. A training and test+validation split of 80:20 resulted in the best performance.

We also tested the effect of data splits on SHRED accuracy. For convention, an *x* : *y* split in data represents a split with *x*% train, *y/*2% validation, and *y/*2% test. We conducted five independent runs each for the following splits {95:5, 90:10, 85:15, 80:20, 70:30, 60:40, 50:50,40:60,30:70} and averaged the train and test MSEs. The data split does not seem to have a significant impact on the accuracy of the reconstruction of the training data. However, increasing the proportion of test data increased the error in the reconstruction of the test dataset. An 80 : 20 split yielded the best results, thus the majority of the experiments conducted for the nematode and the mice used this split.

Applications of SHRED on the nematode *C. elegans* dataset demonstrated the flexibility and adaptability of this neural-network architecture. From only three neurons, SHRED could predict population-wide activity for individual worms and across organisms. We next increased the challenge by leveraging SHRED for a more complex dataset: mouse unit and LFP recordings.

### 3.2 Mouse

The mouse is a popular model organism to study behavior and human diseases [46]. In recent years, we have seen a rapid rise in techniques for large-scale neural recordings during animal behavior, opening the doors to understanding network-level neural codes in the mouse brain [47, 48]. The next dataset we utilized is from the The Visual Coding – Neuropixels project developed by the Allen Brain Observatory (https://observatory.brain-map.org/visualcoding/). The main objective of the Neuropixel project was to study neuronal response to visual stimuli. Mice were presented with various visual cues such as gabors, flashing lights, and drifting gratings. The neural activity —both individual neurons and Local Field Potentials (LFP)— in response to these stimuli were recorded from 6 neuropixel probes inserted throughout the visual cortex.

Each probe contained 374 or 383 channels with each channel containing two bands. The “spike band” measured action potentials of nearby neurons and was recorded at 30 kHz. The “LFP band” measured voltage fluctuations and was recorded at 2.5 kHz. LFPs capture lower frequency voltagefluctuations resulting from transmembrane currents from multiple neurons. Further processing and filtering were conducted to produce the publicly available Neurodata Without Borders (NWB) files. Pupil diameter measurements were also provided for the duration of the recordings. For convention, individual neurons were referred to as “units” in the spiking dataset since it can not be guaranteed that each recording originates from a single cell. Therefore, for the following sections, we also refer to neurons as “units.”

To simplify analysis, we only selected data from a single mouse. A 96-day-old male mouse (ID 756029989) was selected due to the quality of its recordings. Both unit and LFP data were used in the following sections to demonstrate the robustness of SHRED. From reconstructing LFP and average spiking data (Sections 3.2.1 - 3.2.2) to predicting LFP from either an adjacent neurons firing rate (Section 3.2.3) or pupil area (Section 3.2.4), we applied SHRED in a variety of contexts and saw promising results.

#### 3.2.1 SHRED reconstruction of Neural Response to Visual Stimuli

Our first objective was to reconstruct Local Field Potential (LFP) responses to visual stimuli. In this section, we obtained data for the first three presentations of 1) a 2-second 180^*°*^ drifting grating stimulus and 2) a 25-millisecond light flashing stimulus. Visualizations of these stimuli can be found in Figure 7. We selected LFP recordings from both probe A and probe C. Probe A was inserted into the anteromedial (AM) region of the visual cortex and contains a total of 74 channel recordings from the anteromedial area (VISam), cornu ammonis 1 (CA1), dentate gyrus (DG), and the anterior pretectal nucleus (APN). Probe C was inserted into the primary visual cortex (V1) and contains a total of 76 channel recordings from the primary visual cortex (VISp), subiculum (SUB), postsubiculum (POST), superior colliculus (SCig), midbrain (MB), anterior pretectal nucleus (APN), and the posterior limiting nucleus of the thalamus (POL).

**Figure 7.**
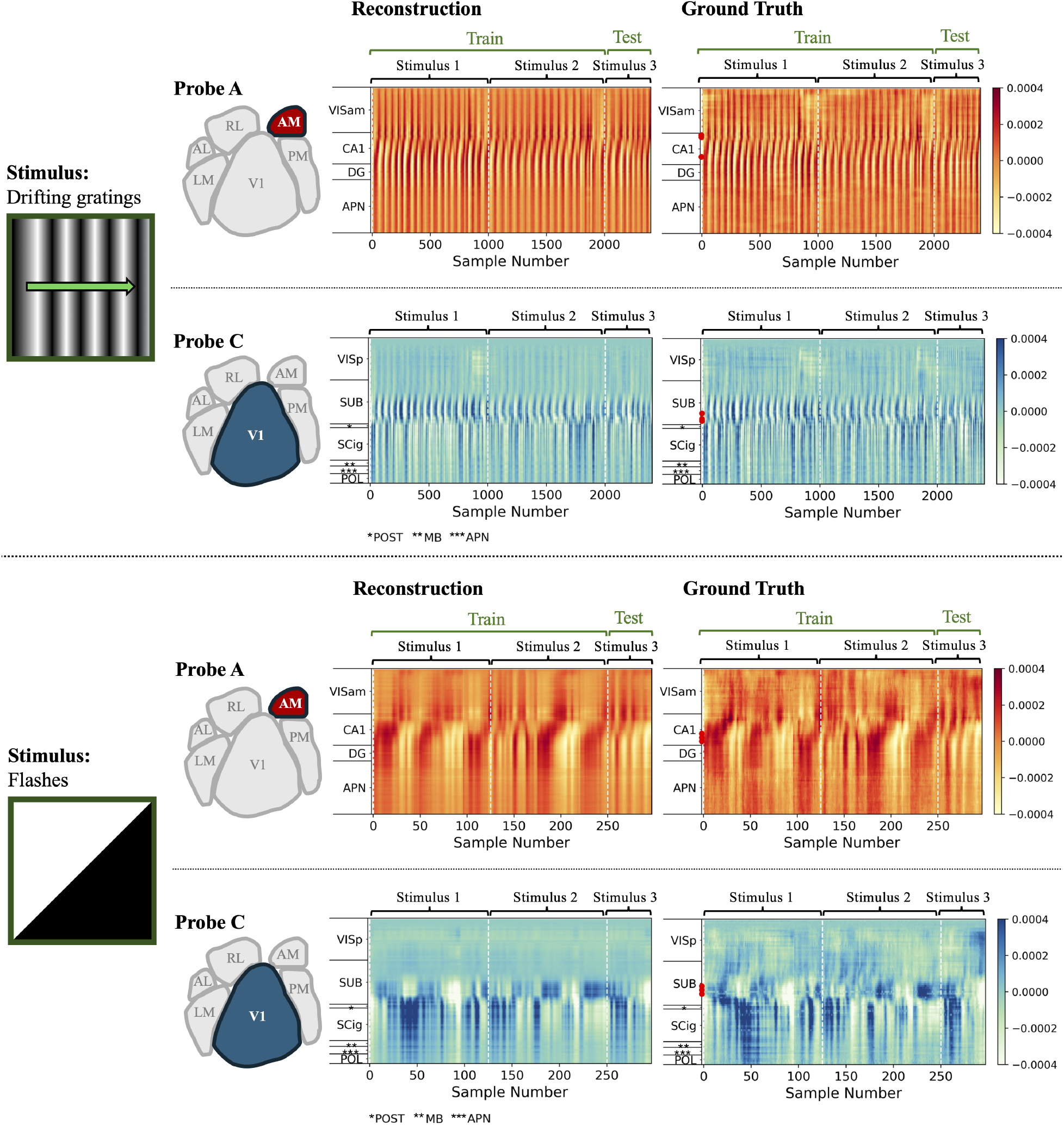
SHRED can accurately reconstruct local field potential in response to a 2s drifting grating and a 25 ms light flashing stimulus. (Stimulus) Local field potential (LFP) recordings were obtained in response to the first three 2-second presentations of a 180^*°*^, 0.8-contrast drifting grating stimulus (top) and 25-millisecond presentations of a 0.8-contrast light flash stimulus (bottom). **(Probe A)** Local field potentials (LFPs) were recorded from the anteromedial (AM) region of the mouse visual cortex. Red dots on the ground truth axis indicate the three neurons selected from the cornu ammonis 1 (CA1) region for training. **(Probe C)** Local field potentials (LFPs) were recorded from the primary visual cortex (V1) of the mouse. Red dots on the ground truth axis indicate the three neurons selected from the high variance subiculum (SUB) region for training. **(Reconstruction)** Recordings from the first two presentations of the stimulus were used for training, while the third presentation served as the test dataset. For both probes, SHRED accurately reconstructed visually evoked LFP responses.

We first reconstructed LFP response to the drifting grating stimulus. The data was downsampled so that each response period contained 1000 timepoints thus were recorded with a sampling rate of 0.5 kHz. The training data was split to include only the first two stimulus presentations. The first half of the third stimulus presentation was set aside as the test dataset, and the last half of the third stimulus presentation was assigned to the validation set. For the drifting grating stimulus, we randomly selected three channels from the cornu ammonis 1 (CA1) region of the hippocampus and trained with a lag of 200 for 200 epochs. For the flash stimulus, we randomly selected three channels from a subset of subiculum (SUB) channels with high variance in activity and trained with a lag of 200 for 200 epochs.

SHRED more accurately reconstructed LFP data from Probe A compared to Probe C. This discrepancy is evident in the carpet plot reconstructions in Figure 7 (top). SHRED struggled to reconstruct activity recorded on Probe C from the primary visual cortex (VISp) region. SHRED was unable to learn the sustained high activity recorded from several channels during the first stimulus presentation. Nevertheless, general patterns in activity are conserved in the reconstruction of both Probe A and Probe C activity. Train and test MSE are quite similar, highlighting the generalizability of SHRED (Table 1(B.i)).

We then reconstructed LFP response to the light flashing stimulus. The data was downsampled so that each response period contained 125 timepoints thus were recorded with a sampling rate of 0.5 kHz. We performed the train-test-val split similar to the drifting grating experiment above. We randomly selected three channels from the cornu ammonis 1 (CA1) region of the hippocampus or from a subset of subiculum (SUB) channels with high variance and trained with a lag of 30 for 200 epochs.

Similarly to the drifting grating stimulus, general patterns in the data are preserved in the flash reconstruction, but SHRED struggles to reconstruct finer details, especially for the Probe C channels in the VISp region. This is reflected in the errors presented in Table 1(B.i). MSE values are slightly higher for the flash stimulus, compared to the drifting grating, most likely due to the smaller dataset size for the shorter duration stimulus. Overall, all reconstructions in this section demonstrate SHRED’s ability to learn larger scale spatio-temporal patterns in population codes. By leveraging these patterns learned, SHRED can reconstruct whole-population activity from only three sensors.

#### 3.2.2 SHRED Reconstruction of Average Unit Responses to Visual Stimuli

SHRED was next utilized to reconstruct averaged unit spike trains in response to all 2-second drifting grating and 25-millisecond flash stimuli. Spike train data is inherently stochastic and consists of discrete, binary events, presenting a significant challenge for many machine learning methods. We averaged spike train data across all presentations of each stimulus type to obtain a more continuous estimate of each unit’s response. Through preliminary filtering of data, we excluded all units with signal to noise ratios *SNR <* 3. For each 2-second presentation of the drifting grating (dg) stimulus, we obtained 2000 time points of spiking data for 243 units. We then averaged this data across presentations to obtain average unit responses. For the 25-millisecond flash stimulus, we obtained 1000 time points of spiking data from 243 units. Note that the sampling rate (SR) differs between the stimuli. Reconstruction of the flash response with the same sampling rate resulted in subpar results due to limitations in data. Therefore, the SR for the flash stimulus was increased. The flash response was also averaged across presentations to obtain the average unit responses. Three units with high variability in activity were randomly selected and the model was trained for 200 epochs.

SHRED more accurately reconstructed the average unit response to the drifting gradient stimulus compared to the flashes. This is most likely due to lower fluctuations in individual spiking activity across time for the DG stimulus compared to the flashes (Figure 8). We observe lower fluctuations most likely due to two main reasons: 1) the drifting grating unit response is sampled with a lower sampling rate and 2) responses to drifting gratings of different orientations, speed, and contrasts are averaged together, smoothing out the unit’s response. For example, consider a unit with a receptive field sensitive to light located in the bottom left-hand corner of the screen. When gratings are presented at different orientation and at different speeds, it is plausible that the receptive field will always be lit in at least one of these presentations. Therefore, averaging the data may result in a smoother activity trace, setting up an easier reconstruction task for SHRED.

**Figure 8.**
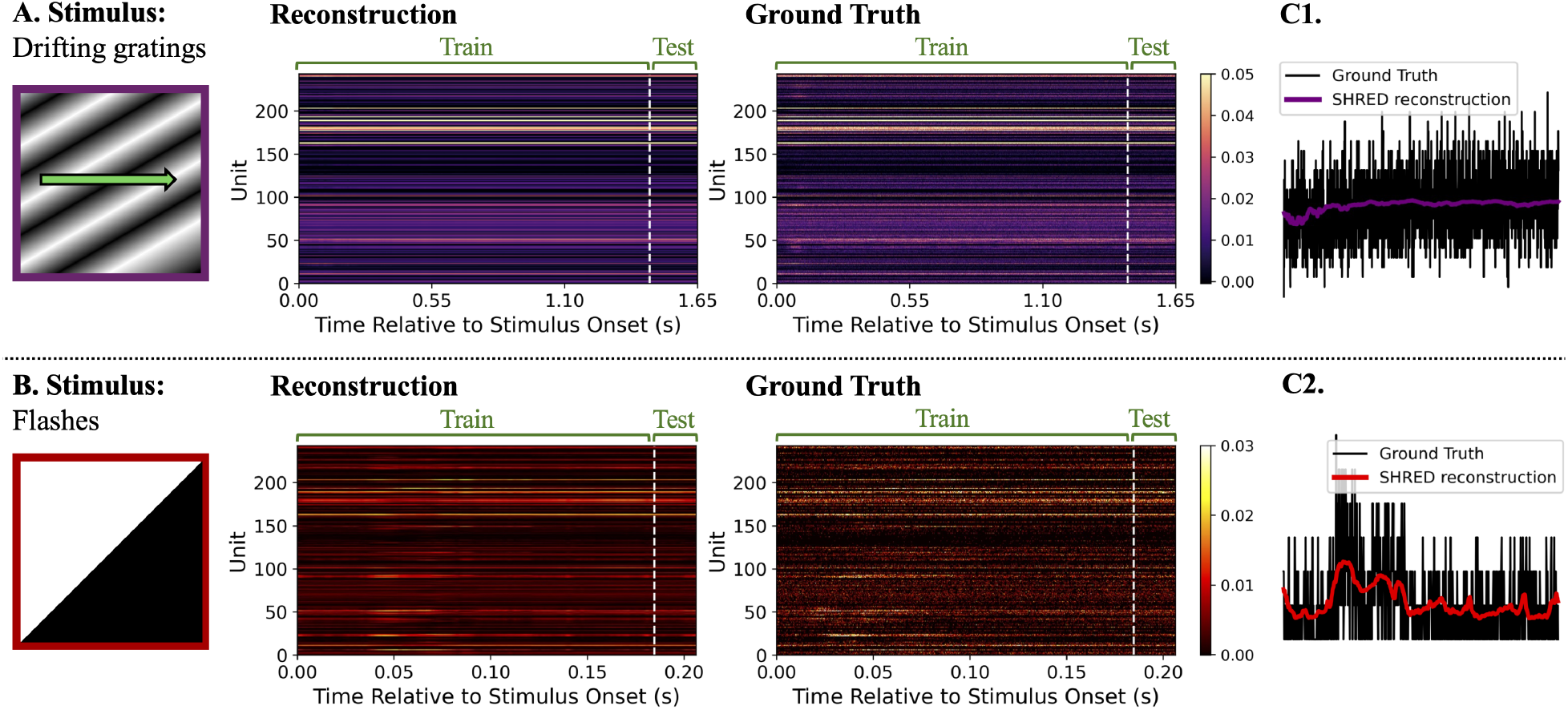
SHRED accurately reconstructs and filters high frequency recordings of averaged unit activity. **(A)** SHRED reconstruction of average unit activity for all 2-s trials with drifting grating stimuli. **(B)** SHRED reconstruction of average unit activity for all 25-ms trials with light flashing stimuli. **(C1-C2)** Average activity (black) and SHRED reconstruction (purple/red) of a randomly selected neuron. For both stimuli, averaged unit recordings contain high frequency fluctuations which are filtered out in the SHRED reconstruction. Overall trends in activity are well-preserved.

While the reconstruction errors are higher for the flash stimulus (Table 1(B.ii)) the example traces of unit 150 in Figure 8 C1-C2 show that general activity patterns are preserved. While individual spikes are not reconstructed, SHRED learns the time course of when the unit is firing more frequently. From this result, we hypothesized that the model is learning the overall firing rates rather than the timing of the individual spikes. Therefore in the following section, we focus on reconstruction of LFP from firing rate.

#### 3.2.3 SHRED Reconstruction of Local Field Potential from Individual Neuron Firing Rate

In Section 3.2.2, we observed that SHRED learns overall firing rates of individual units. Therefore, we then hypothesized that SHRED could reconstruct local field potential (LFP) from firing rate data. We obtained unit recordings from probe A in response to the first 2-second, 180^*°*^, 0.8 contrast drifting grating stimulus. Out of the units with firing rate *fr >* 10, we selected the first unit for this analysis. We then obtained the LFP recording from the same channel, which reflects voltage fluctuations in the extracellular space near the selected unit.

We then obtained the firing rate of the selected unit from the spike train. Firing rate (*v*) of a neuron is defined as

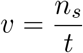

where *n*_*s*_ is the number of spikes in time *t*. To obtain an accurate measurement of the spike rate, we used a one-dimensional gaussian filter. The sampling rate (*sr*) for the unit data was down sampled to around 1.25 kHz. The standard deviation for the gaussian was set to

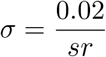

This means that we smoothed the spike train over a 20 millisecond window. After applying the filter, we multiplied the filtered trace by the sampling rate to obtain the firing rate in spikes/s. The smoothed firing rate is shown in Figure 9 A. We included 2002 time points in the training, 250 in the validation, and 250 in the testing dataset. SHRED was trained to predict the LFP and reconstruct the firing rate.

**Figure 9.**
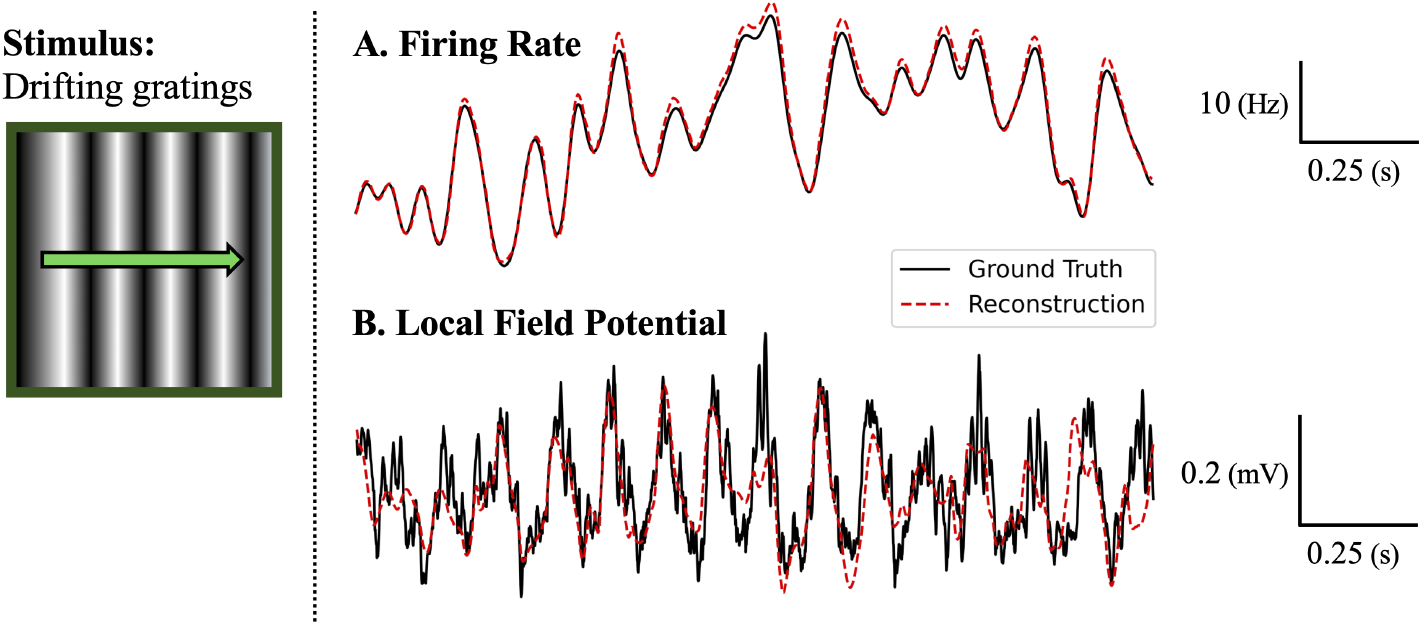
SHRED can reconstruct Local Field Potential (LFP) from firing rate of nearby neuron. **(A)** Firing rate of a neuron in the visual cortex located near the probe inserted in the anteromedial (AM) region served as the input to the SHRED model. **(B)** The SHRED model was trained to reconstruct LFP recording closest to the neuron selected as input. The model reconstruction (red) closely follows the general patterns of the LFP recoding (black).

From just the neuron firing rate, SHRED can predict local field potential (Table 1(B.iii)). The SHRED reconstruction of the entire dataset shown in Figure 9 closely follows the data. While there are some deviations between the predicted and the true LFP, overall activity trends are well-preserved. It is evident that the firing rate is dynamically coupled to the LFP, making this a suitable setting for applying SHRED. Based on the conclusions of this experiment, we hypothesized that any dataset that is dynamically coupled to neural recordings should be sufficient to reconstruct population-wide neural activity using SHRED. Therefore, in the next experiment outlined in Section 3.2.4, we reconstructed LFP data from just the pupil area recording.

#### 3.2.4 SHRED Reconstruction of Local Field Potential from Pupil Area Response to a Visual Stimulus

In theory, pupil area should be dynamically coupled to the local field potential in response to a visual stimulus [49]. Therefore, in this experiment, we aimed to reconstruct LFP recordings from just the pupil area measurements. Probe A LFP and pupil area recordings were obtained from the first presentation of the 2-second, 180^*°*^, 0.8 contrast drifting grating stimulus. To account for the different sampling rates, we applied a cubic spline to the pupil area measurements. Thus pupil recordings were obtained for the same timepoints as the LFP recordings. We split the data into 75% train, 12.5% test, and 12.5% validation. With a lag of 50, pupil area recordings served as the input to the model which was trained to predict the LFP and reconstruct the pupil area recordings. Although this was a very difficult task at hand, SHRED performed surprisingly well. The reconstruction shown in Figure 10 deviates from the ground truth significantly, but the peaks and troughs are still maintained. Train and test MSE are presented in Table 1 (B.iv). While there is still more work needed to improve the prediction accuracy, this baseline result highlights the versatility and robustness of the SHRED architecture.

**Figure 10.**
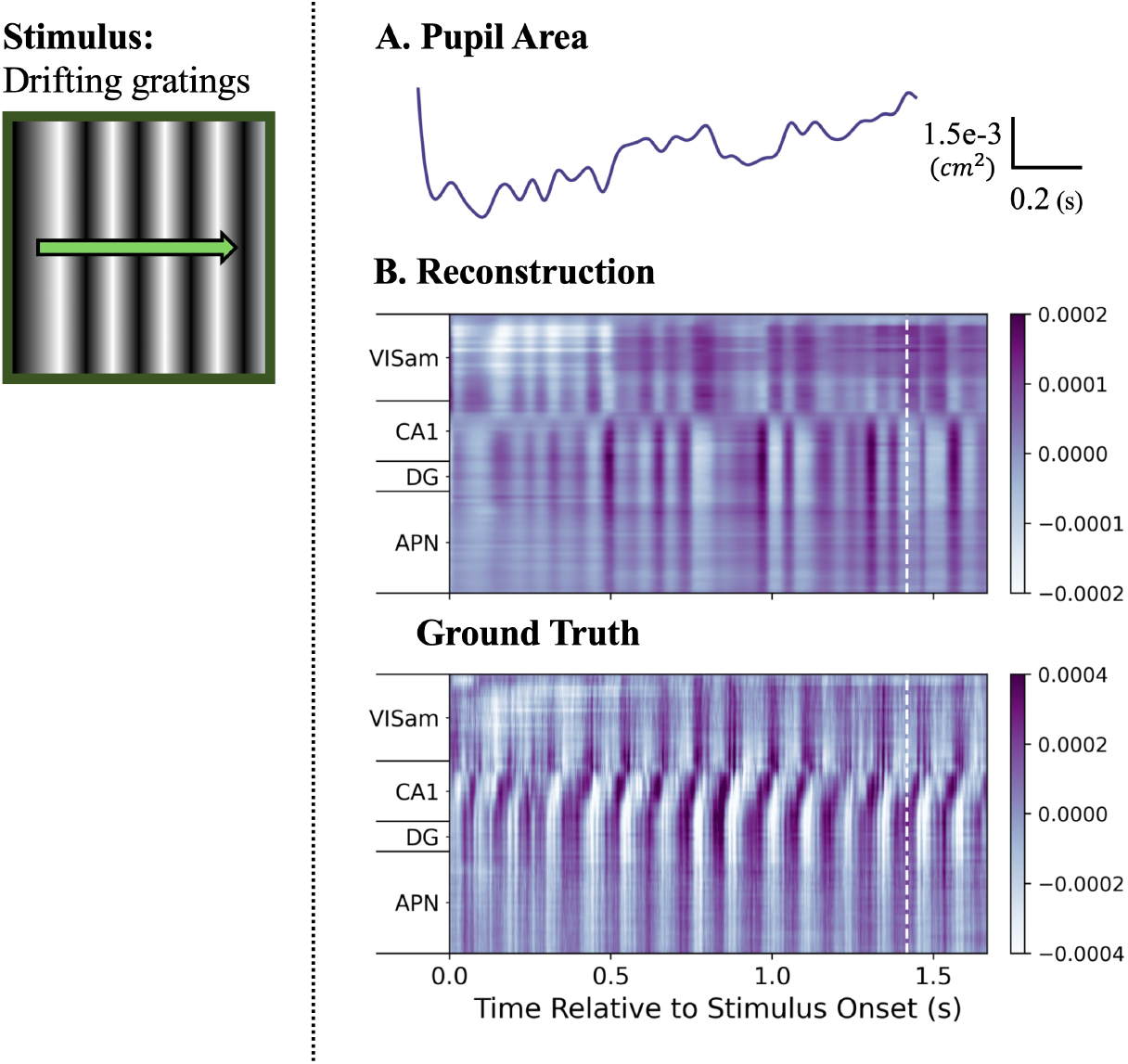
SHRED has potential to reconstruct Local Field Potential (LFP) activity from pupil area. **(A)** Pupil area in response to a 2-s drifting grating stimulus served as the input to the SHRED model. **(B)** The SHRED model was trained to reconstruct LFP activity from a probe inserted in the anteromedial (AM) region of the mouse visual cortex. The reconstruction successfully captures coarse fluctuations in spatial and temporal activity.

Applications of SHRED to the mouse dataset highlighted it’s diverse usage. Not only is SHRED capable of reconstructing population neural activity from a few sensors, it is also capable of reconstructing population dynamics from any coupled measurements. By learning the patterns and connections in these datasets, SHRED was capable of predicting general brain activity from unit firing rate and pupil area measurements. To further explore the versatility of SHRED, we next reconstructed a video of the zebrafish forebrain activity.

### 3.3 Zebra Fish

The zebrafish, a small freshwater species in the minnow family, is a widely used model organism for studying vertebrate development and, more recently, systems neuroscience [50, 51]. Zebrafish offer several advantages in research, one of which is their transparency during early developmental stages, enabling clear and accessible imaging [50,52]. Recent studies have focused on understanding neuronal ensembles code for sensorimotor pathways [3, 6, 53–55]. With improvements in imaging technologies, high-speed light-sheet microscopy can now be implemented to simultaneously record from almost all neurons in the zebrafish. This large scale recording allows for complex analysis of activity patterns in the whole brain [56].

For our third model system, we looked at a video of forebrain activity of a zebra fish obtained using high-speed light-sheet microscopy. This video was made publicly available by [56]. Neural projections were genetically encoded using a calcium indicator GCaMP5G. The 1041.859-second video contained 750 frames indicating a frame rate of approximately 0.72 Hz. We then converted the video to grayscale and cropped each frame to include 472 *×* 600 pixels thus a total of 283,200 pixels.

We aimed to reconstruct this video from the timeseries of just 250 pixels. We split the data to include 700 time points in the training set, 25 time points in the testing set, and 25 time points in the validation set and trained the model for 200 epochs.

SHRED struggles to reconstruct individual changes in pixels, but captures flares of activity with great accuracy (Figure 11). Train and test MSE are presented in Table 1 (C.i). The overall shape of the forebrain was preserved and regions with increased activity was reflected in the reconstruction. In this section, we showed that SHRED can be used to reconstruct videos. SHRED is versatile and can be applied to a variety of complex datasets with minimal hyperparameter tuning.

**Figure 11.**
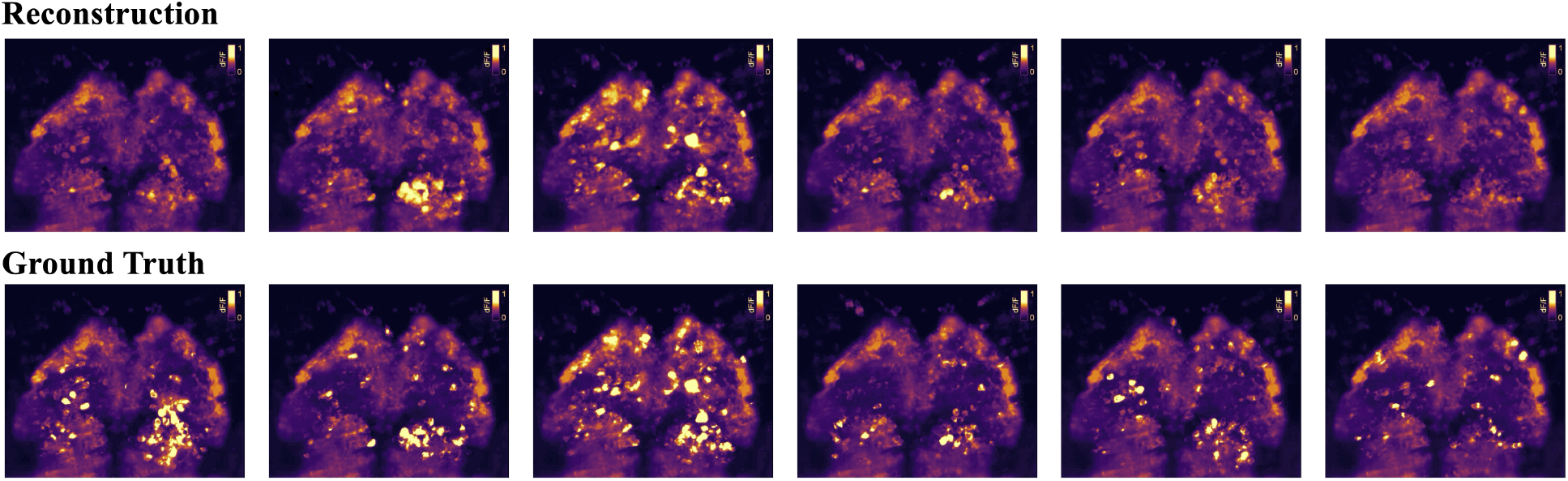
SHRED accurately reconstructed a video of zebra fish forebrain activity. SHRED reconstruction of 6 time points (top) follows general trends in activity of the original (bottom).

### 3.4 Human Locomotion

In the final data set for consideration, we highlight the recent work of Ebers et al [15] where SHRED was used for mapping temporal sensor measurements to behavior. Specifically, SHRED is used with mobile sensing in order to monitor human locomotion and biomechanics. The tracking and analysis of human motion is critical for many tasks, including the monitoring of disease progression, guiding rehabilitation treatment, evaluating sports performance, and informing assistive device design. Tracking motion in general is an expensive endeavor requiring a myriad of sensors. However, with SHRED one can replace the array of sensors monitoring motion with a single smart watch and its accelerometer. The data set for this mobile SHRED study [15] is an open-source dataset that captures nondisabled human biomechanics during steady, rhythmic walking; such an approach can be generalized to other motion tracking applications like robotic manipulation or computer animation. We use the time series of joint kinematics from 12 nondisabled adults (six female/six male; age 23.9 *±* 1.8 years; height = 1.69 *±* 0.10m; mass = 66.5 *±* 11.7 kg) [57]. Participants walked at a self-selected speed (speed = 1.36 *±* 0.11 m/s) on a split-belt instrumented treadmill (Bertec Corp., Columbus, USA) for six minutes. While kinematics can be captured with multiple modalities, gold-standard marker data was recorded using a 10-camera optical motion capture system. Kinematics were estimated from marker data using the Inverse Kinematics algorithm in OpenSim 3.3 with a dynamically-constrained 19 degree-of-freedom skeletal model scaled to each participant [58, 59]. The SHRED work evaluated the 18 kinematic features previously calculated in the open-source dataset, which included three translational degrees of freedom at the pelvis, pelvis tilt/list/rotation, lumbar extension/bending, bilateral hip flexion/adduction/rotation, bilateral knee flexion, and bilateral ankle dorsiflexion. Kinematics were low-pass filtered at 6Hz using a fourth-order Butterworth filter [57]. An important distinction with this human biomechanics data, as opposed to the neural data used earlier, is that marker data recorded during motion capture is in a global coordinate frame, but through the standard procedures for marker data processing are transformed into kinematics in a local coordinate frame relative to the human subject [60].

For the population-based biomechanics models, a reconstruction mapping was trained using 11 of the 12 participants’ kinematic data, for a one-subject hold-out test using each modeling paradigm. Two combinations of dynamic trajectory inputs were tested: (1) three non-randomly chosen kinematic features (right hip flexion angle, right knee flexion angle, right ankle dorsiflexion angle), and (2) one non-randomly chosen kinematic feature (right ankle dorsiflexion angle).

Figure 12 summarizes the performance of SHRED relative to the SDN and linear models for reconstructing human motion within a population. SHRED far outperforms the SDN and linear model regardless of number or choice of input sensors. Using three mobile sensors, SHRED has errors less than 0.098 *±* 0.062 degrees, which was on average 3.3x more accurate than the SDN and 9.6x more accurate than the linear model. Using a single mobile sensor, SHRED has errors less than 0.121 *±* 0.073 degrees, which was on average 9.8x more accurate than the SDN and 14.8x more accurate than the linear model. These results demonstrate that SHRED can enable rapid generalization (parameterization of dynamics) for data outside the training set, which may be especially useful in human biomechanics where population-level models can be used to estimate parameters for a new individual (*e*.*g*., a new patient visits a clinic for the first time).

**Figure 12.**
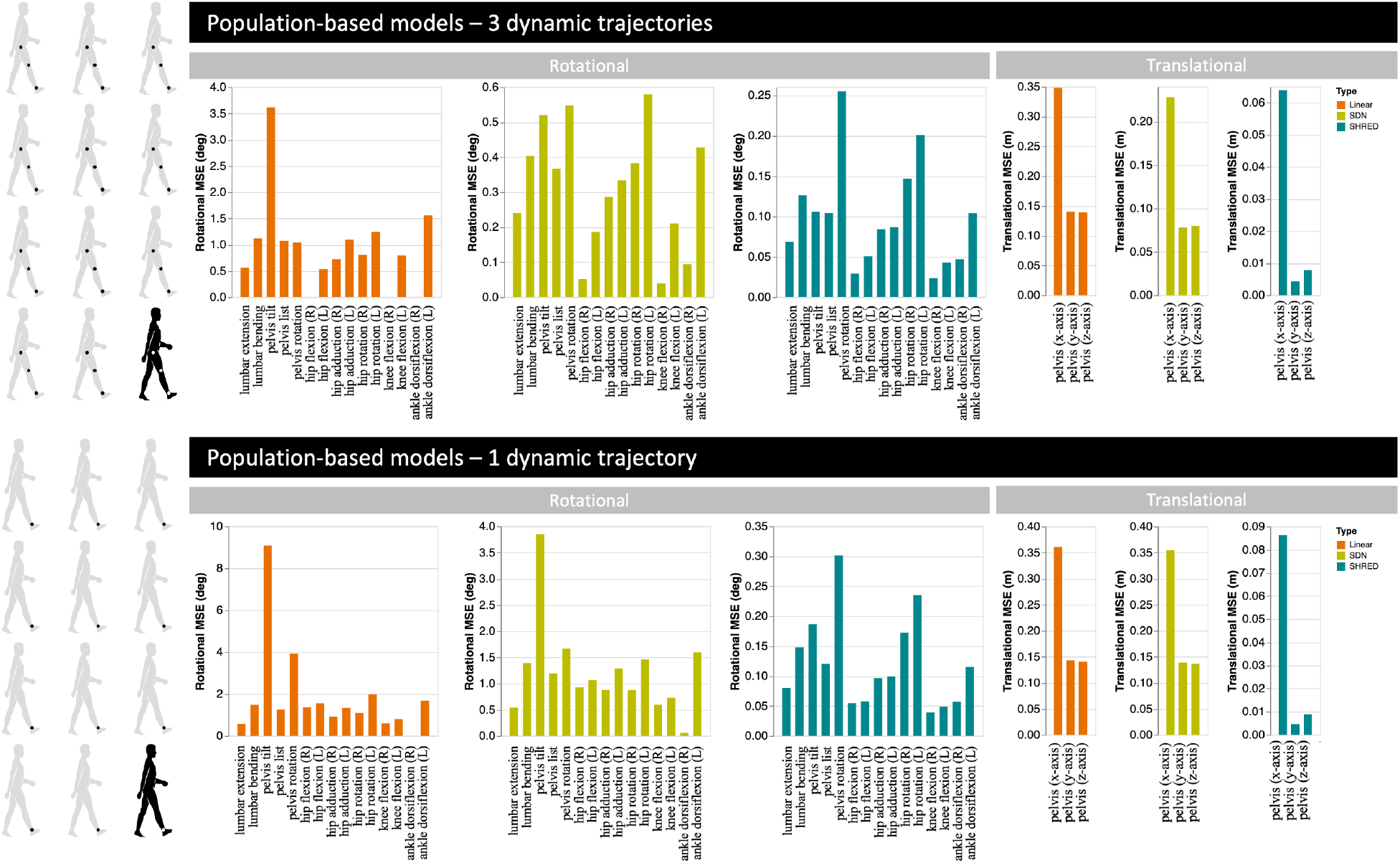
Visualization of population-based results from Ebers et al [15] for reconstructing human biomechanics [57]. The population models trained using 11 of the 12 individuals’ data and were evaluated by reconstructing the held-out subject’s full measurement data from their sparse input sensor(s); performance was calculated by using mean-squared error (MSE). Rotational and translational kinematic states are displayed separately, as rotational states use units of degrees while translational states use units of meters. Three modeling paradigms were trained to learn the mapping: linear regression (orange), a shallow decoder neural network (green), and a shallow recurrent neural network (blue). Note the independent y-axes and respective scales for each graph. (Top) Three dynamic trajectories were purposefully chosen as sensor inputs to the reconstruction mapping: right hip flexion angle (degrees), right knee flexion angle (degrees), and right ankle dorsiflexion angle (degrees). (Bottom) One dynamic trajectory was purposefully chosen as sensor inputs to the reconstruction mapping: right ankle dorsiflexion angle (degrees). [Reprinted with permission from Ebers et al [15]]

## 4 Conclusions and Discussions

Modern neuroscience and behavioral sciences are now supplemented with exceptional data sets which measure a diversity of quantities, from individual neurons to whole brain activity to behavioral outcomes. Such emerging data sets, which are now routinely in the open source domain, offer significant scientific opportunities for learning and discovery in the biological sciences. Specifically, there are modern data-driven mathematical architectures and algorithms capable of extracting interpretable generative models from the diversity of data collected. Moreover, it is ideal if cheap and non-invasive sensing can be used instead of expensive and invasive sensors. The SHRED architecture introduced in this work provides a viable pathway towards using a highly limited number of sensors for reconstructing the full high-dimensional state-space of either neural activity or behavioral responses. This is demonstrated on a number of toy models in biology and on behavioral data for locomotion.

SHRED is an exceptionally flexible architecture, being now used for many engineering and physics applications [14–18]. Moreover, it has been demonstrated to have a globally convex landscape which allows for rapid training with little or no hyper parameter tuning [18]. SHRED is ideal for applications in the biological sciences which requires flexibility as many applications are in low-data limits with large measurement noise. SHRED can operate in this difficult data limit as easily as in the high-data limit. And importantly, because of the globally convex landscape and ability to train with compressed data [14, 17], training SHRED with neural data can be accomplished on laptop-level computing, even for exceptionally large data sets.

SHRED as demonstrated in this work to have the capability of mapping limited neural or behavioral recordings accurately to full state-space measurements. The success of the method allows for deployment in practice to new patients or model organisms. For both the *C. elegans* and biolocomtion examples, the ability of SHRED to be deployed in a parametric fashion affords many opportunities in clinical settings where non-invasive measurements can be used to approximate difficult or invasive measurements. In future work, learning generative models of the dynamics will be explored with the goal of extracting even more meaningful insights into biological activity [18]. Thus SHRED overall is an emerging deep learning algorithm which is well suited for diverse biological data.

## 5 Code and Data

All code and data are publicly available. Figures and experiments can be reproduced from the following GitHub:

https://github.com/amysrude/neuralSHRED.

The data sets used in the analyses are from the following:

*C. elegans*: https://osf.io/2395t/

Mouse: https://allensdk.readthedocs.io/en/latest/visual_coding_neuropixels.html

Zebrafish: https://www.nature.com/articles/nmeth.2434 biolocomotion: https://github.com/meganebers/mobileSHRED and [57–59]

## Acknowledgements

We wish to acknowledge the support of the National Science Foundation AI Institute in Dynamic Systems grant 2112085 and support from the Air Force Office of Scientific Research (FA9550-24-1-0141).

## Notes

https://osf.io/2395t/

https://allensdk.readthedocs.io/en/latest/visual_coding_neuropixels.html

https://www.nature.com/articles/nmeth.2434

https://github.com/meganebers/mobileSHRED

### Competing Interest Statement

The authors have declared no competing interest.

